# Lysosomal TMEM106B interacts with galactosylceramidase to regulate myelin lipid metabolism

**DOI:** 10.1101/2023.09.14.557804

**Authors:** Hideyuki Takahashi, Azucena Perez-Canamas, Hongping Ye, Xianlin Han, Stephen M. Strittmatter

## Abstract

TMEM106B is an endolysosomal transmembrane protein not only associated with multiple neurological disorders including frontotemporal dementia, Alzheimer’s disease, and hypomyelinating leukodystrophy but also potentially involved in COVID-19. Additionally, recent studies have identified amyloid fibrils of C-terminal TMEM106B in both aged healthy and neurodegenerative brains. However, so far little is known about physiological functions of TMEM106B in the endolysosome and how TMEM106B is involved in a wide range of human conditions at molecular levels. Here, we performed lipidomic analysis of the brain of TMEM106B-deficient mice. We found that TMEM106B deficiency significantly decreases levels of two major classes of myelin lipids, galactosylceramide and its sulfated derivative sulfatide. Subsequent co-immunoprecipitation assay showed that TMEM106B physically interacts with galactosylceramidase. We also found that galactosyceramidase activity was significantly increased in TMEM106B-deficient brains. Thus, our results reveal a novel function of TMEM106B interacting with galactosyceramidase to regulate myelin lipid metabolism and have implications for TMEM106B-associated diseases.

## Introduction

*TMEM106B*, which encodes TMEM106B, a highly glycosylated type II transmembrane protein, was initially identified as a risk modifier of frontotemporal lobar degeneration (FTLD) especially in patients with mutations in the *GRN* (FTLD-*GRN*) (Feng et al., 2021; Perneel and Rademakers, 2022; Van Deerlin et al., 2010). Subsequently, *TMEM106B* has also been linked to many other neurodegenerative diseases, such as Alzheimer’s disease (AD) (Bellenguez et al., 2022; Hu et al., 2021; Wightman et al., 2021), chronic traumatic encephalopathy (Cherry et al., 2018), and limbic-predominant age-related TDP-43 encephalopathy (Nelson et al., 2019). In addition, a *de novo* mutation in *TMEM106B* was found in patients with hypomyelinating leukodystrophy (HLD), a neurodevelopmental disorder characterized by myelin deficits (Simons et al., 2017). Outside the brain, previous studies using an in vivo genetic screen have identified *TMEM106B* as a primary robust driver of lung cancer metastasis (Grzeskowiak et al., 2018; Kundu et al., 2018). Furthermore, three different studies using genome-wide CRISPR screens independently identified TMEM106B as a host factor in certain cell lines for SARS-CoV-2, a strain of coronavirus that caused coronavirus disease 2019 (COVID-19) (Baggen et al., 2021; Schneider et al., 2021; Wang et al., 2021).

While *TMEM106B* has been shown to be associated with a great variety of human conditions, little is known about molecular mechanism by which TMEM106B is involved in those conditions. TMEM106B is localized at the endolysosome and thought to regulate several aspects of lysosomal function, but so far, a limited number of the interacting proteins have been reported (Feng *et al*., 2021; Perneel and Rademakers, 2022). Recent studies using cryo-electron microscopy have shown that truncated C-terminal TMEM106B forms amyloid fibrils in the brain (Chang et al., 2022; Fan et al., 2022; Jiang et al., 2022; Schweighauser et al., 2022; Takahashi and Strittmatter, 2022). However, the TMEM106B fibrils have been found not only in the brains of people with neurodegenerative diseases but also in neurologically normal ageing brains. Additionally, a disease-protective variant of TMEM106B also forms the amyloid fibrils. It thus remains unclear whether the amyloid fibrils are involved in the progression of TMEM106B-associated neurological disorders.

Dysregulation of lipid metabolism has been reported in the brains of FTLD-*GRN* patients as well as mouse models of FTLD-*GRN* (Boland et al., 2022; Evers et al., 2017; Logan et al., 2021; Marian et al., 2023). Given the role of *TMEM106B* in FTLD-*GRN*, these findings raise a possibility that TMEM106B also regulates lipid metabolism under physiological and/or pathological conditions. Here, we performed lipidomic analysis of the brain of TMEM106B-deficient mice. We found that levels of two major classes of myelin lipids galactosylceramide (GalCer) and sulfatide (ST) are significantly reduced in TMEM106B-deficient brains. Subsequent unbiased proteomics and co-immunoprecipitation (co-IP) assay showed that TMEM106B physically interacts with galactosylceramidase (GALC), a lysosomal enzyme that hydrolyzes GalCer. We also found that GALC activity was significantly increased in TMEM106B-deficient brains. These results suggest that TMEM106B interacts with GALC to regulate myelin lipid metabolism particularly GalCer and ST levels. Thus, GALC inhibition may be a therapeutic target for TMEM106B-associated diseases.

## Results

### Levels of GalCer and ST are decreased in TMEM106B-deficient brains

Given the potential roles of TMEM106B in regulating lipid metabolism, we performed multidimensional mass spectrometry-based shotgun lipidomic analysis (Han, 2016) using the brain of 12-month-old TMEM106B-deficient mice and wild-type (WT) littermates. Lipid classes analyzed in our lipidomics include phosphatidylcholine (PC), phosphatidylethanolamine (PE), phosphatidylserine (PS), phosphatidylglycerol (PG), phosphatidylinositol (PI), phosphatidic acid (PA), cardiolipin (CL), lyso-phosphatidylcholine (LPC), lyso-phosphatidylethanolamine (LPE), lyso-cardiolipin (LCL), sphingomyelin (SM), ceramide (CER), sulfatide (ST), cerebroside (CBS), free cholesterol (CHL), cholesterol esters (CE), bis(monoacylglycerol)phosphate (BMP), and the total number of lipid species analyzed is 294 (**Figure 1A**). Strikingly, we found that levels of multiple species of CBS and ST were significantly decreased in TMEM106B-deficient brains (**Figure 1B and C**), although TMEM106B deficiency affected levels of several species of other lipid classes such as PA, PE, and PC as well (**Figure 1C**). Importantly, total levels of ST, CBS, and PA were also found to be decreased in TMEM106B-deficient brains (**Figure 1D**).

**Figure 1:**
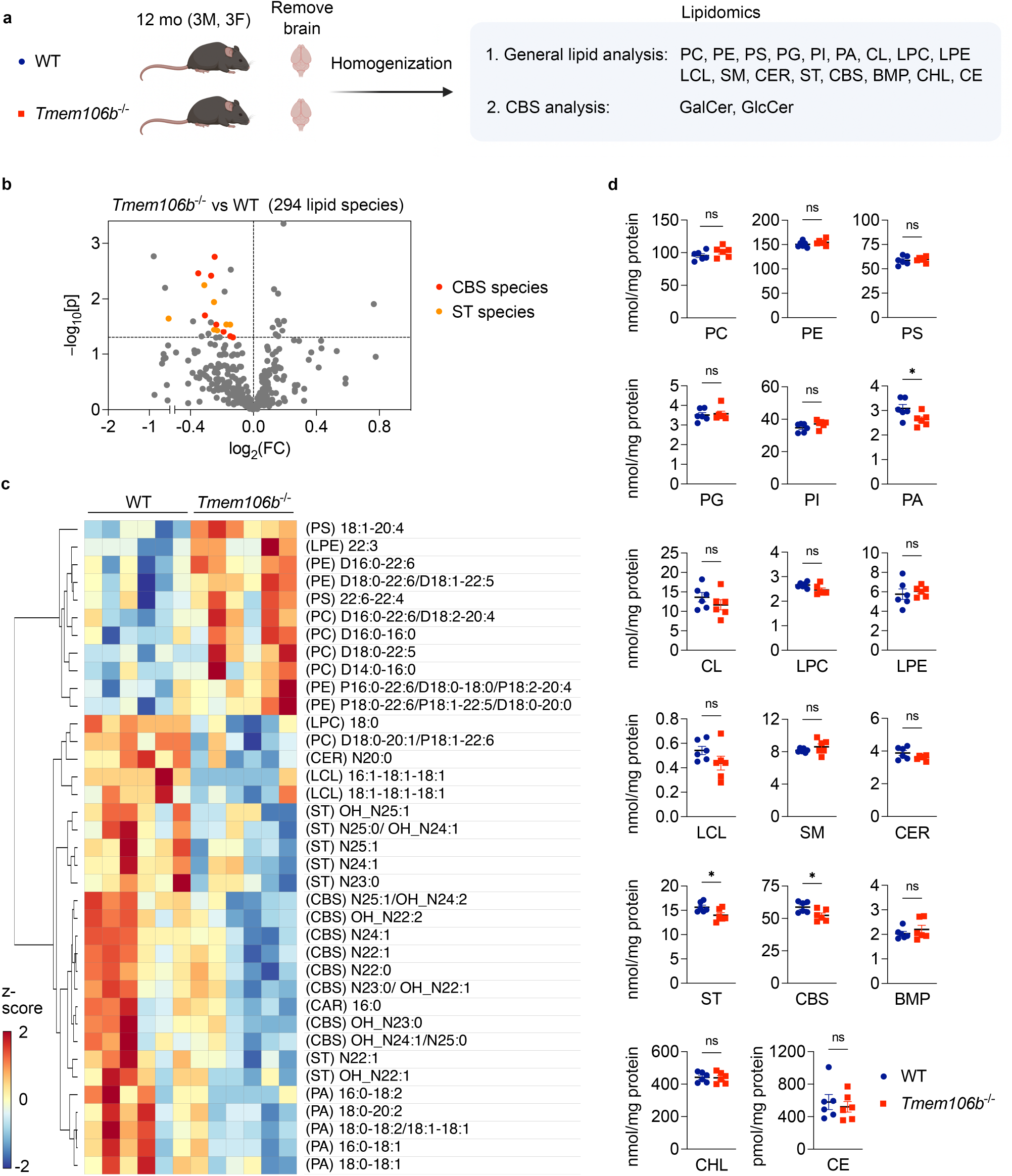
Cerebroside and sulfatide are decreased in TMEM106B-decifient brains. **a)** Diagram showing the experimental procedures of lipidomic analysis using the brain of 12-month-old WT and *Tmem106b*^-/-^ mice. n = 6 (3 males and 3 females) per genotype. First, targeted shotgun lipidomics were performed to analyze 17 general lipid classes; phosphatidylcholine (PC), phosphatidylethanolamine (PE), phosphatidylserine (PS), phosphatidylglycerol (PG), phosphatidylinositol (PI), phosphatidic acid (PA), cardiolipin (CL), lyso-phosphatidylcholine (LPC), lyso-phosphatidylethanolamine (LPE), lyso-cardiolipin (LCL), sphingomyelin (SM), ceramide (CER), sulfatide (ST), cerebroside (CBS), free cholesterol (CHL), cholesterol esters (CE), bis(monoacylglycerol)phosphate (BMP). The total number of lipid species analyzed in this study is 294. SFC-MS/MS was then performed to analyze galactosylceramide (GalCer) and glucosylceramide (GlcCer) species separately. The diagram was created with BioRender.com. **b)** Volcano plot showing all 294 molecular lipid species. CBS and ST species (p < 0.05) are highlighted in red and orange, respectively. The horizontal line indicates p = 0.05. P values were calculated using two-tailed unpaired t-test. **c)** Heatmap of all lipid species significantly affected by TMEM106B deficiency at p < 0.05 cutoff. d) Scatter plots of the sum of all species within a lipid class for each class in WT versus *Tmem106b*^-/-^. Mean ± SEM, n = 6 (3 males and 3 females) per genotype. *p < 0.05; Two-tailed unpaired t-test.

CBS comprises galactosylceramide (GalCer) and glucosylceramide (GlcCer). To determine levels of GalCer and GlcCer species separately, we next performed supercritical fluid chromatography-tandem mass spectrometry (SFC-MS/MS) using the same brain homogenates and found that levels of multiple species of GalCer were significantly decreased in TMEM106B-deficient brains while none of GlcCer species was significantly altered (**Figure 1A and 2A and B**). TMEM106B deficiency also significantly decreased total levels of GalCer but not GlcCer in the brain (**Figure 2C and D**). A decrease in GalCer and its sulfated derivative ST in TMEM106B-deficient brains was of particular interest as GalCer and ST are two major classes of myelin lipids (Marcus and Popko, 2002) and as *TMEM106B* is an HLD gene. Note that TMEM106B deficiency however had no effects on levels of CER and SM (**Figure 1D**), revealing a specific effect of TMEM106B deficiency on GalCer and ST levels. Together, our lipidomic data suggest that TMEM106B deficiency affects myelin lipid metabolism by decreasing levels of GalCer and ST in the brain.

**Figure 2:**
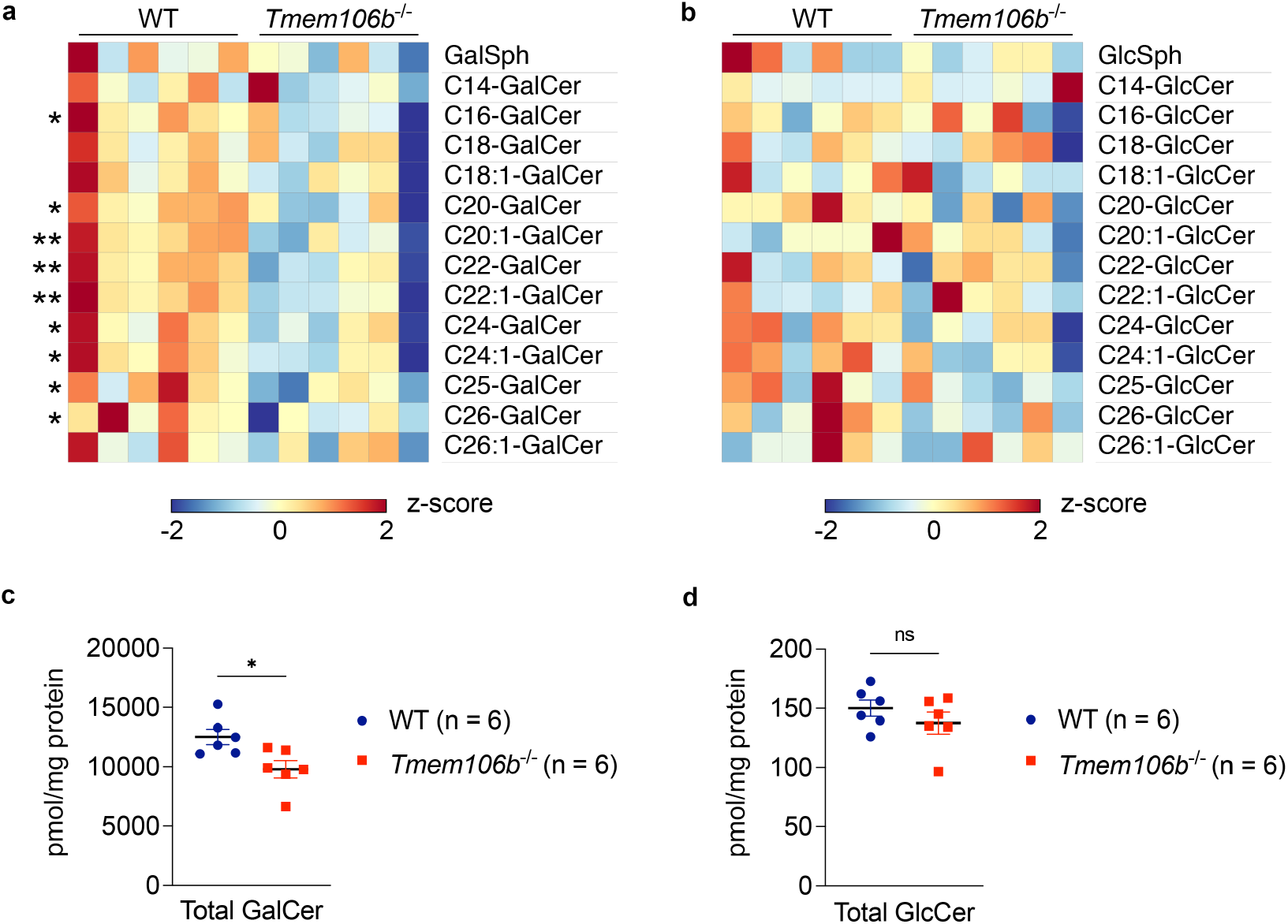
GalCer but not GlcCer is decreased in TMEM106B-deficient brains. **a)** Heatmap showing levels of GalCer species in the brain of WT and *Tmem106b*^−/−^ mice at 12 months of age. n = 6 mice (3 males and 3 females) per genotype. *p < 0.05, **p < 0.01; Two-tailed unpaired t-test. **b)** Heatmap showing levels of GlcCer species in the brain of WT and *Tmem106b*^−/−^ mice at 12 months of age. n = 6 mice (3 males and 3 females) per genotype. None of species are significantly affected by two-tailed unpaired t-test. **c)** Total GalCer levels (sum of all species) in the brain of WT and *Tmem106b*^−/−^ mice at 12 months of age. Mean ± SEM, n = 6 mice (3 males and 3 females) per genotype. *p < 0.05; Two-tailed unpaired t-test. **d)** Total GlcCer levels (sum of all species) in the brain of WT and *Tmem106b*^−/−^ mice at 12 months of age. Mean ± SEM, n = 6 mice (3 males and 3 females) per genotype. Two genotypes were not significantly different by two-tailed unpaired t-test.

### TMEM106B physically interacts with GALC and regulates GALC activity

To examine the mechanisms by which TMEM106B deficiency affects myelin lipid metabolism, we next sought to identify interaction partners of TMEM106B in the brain by using immunoprecitation (IP) and liquid chromatography tandem mass spectrometry (LC-MS/MS) protein identification procedure (**Figure 3A**). We confirmed that our anti-TMEM106B antibody cross-linked to protein A Sepharose beads reproducibly immunoprecipitated endogenous TMEM106B from mouse brains (**Figure 3B**).

**Figure 3:**
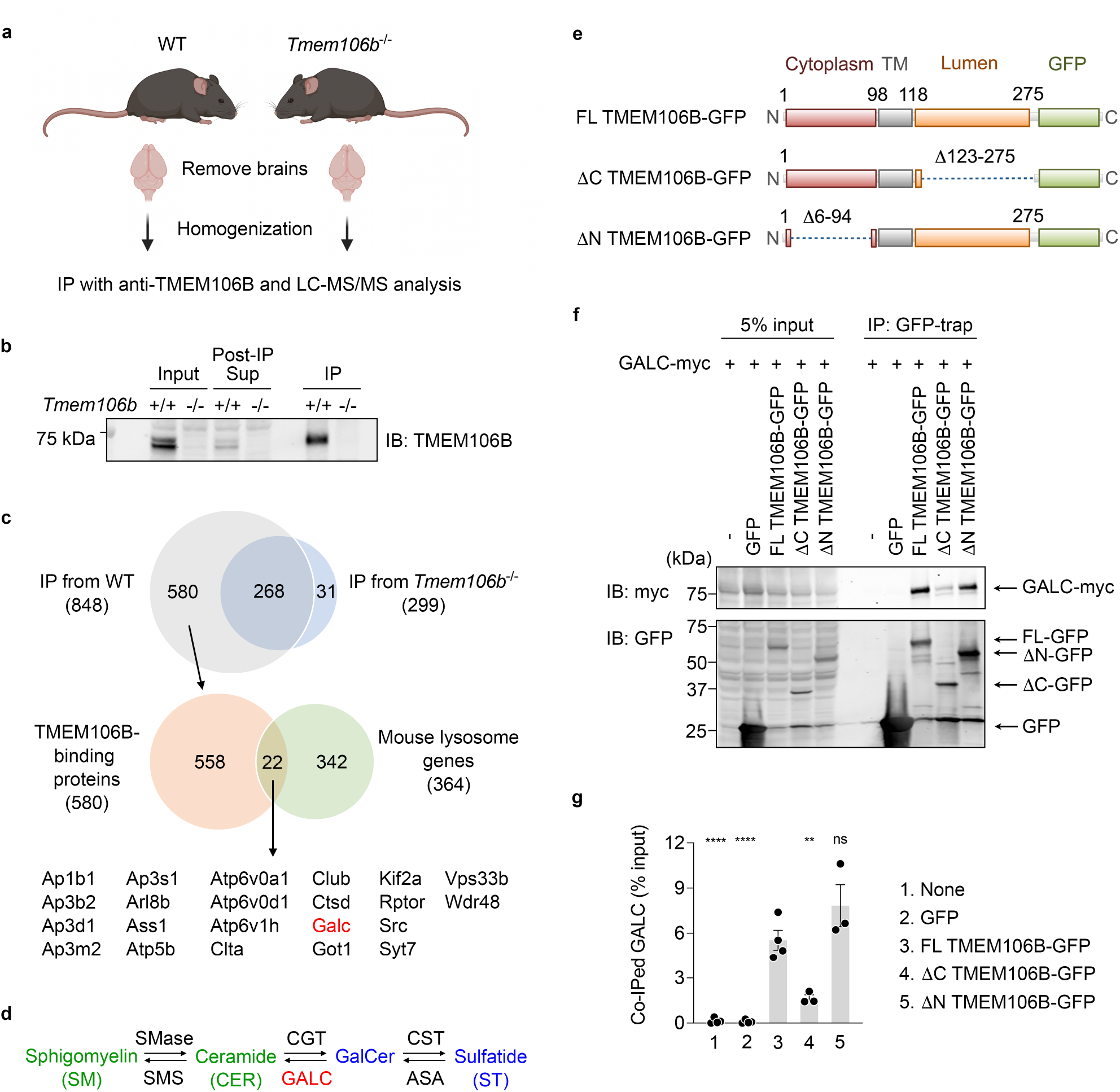
TMEM106B physically interacts with GALC via its lumenal region. **a)** Diagram showing experimental procedures of IP and LS-MS/MS analysis using mouse brains. The diagram was created with BioRender.com. **b)** Representative blots to validate immunoprecipitation of endogenous TMEM106B from mouse brains using anti-TMEM106B antibody linked to protein A Sepharose. **c)** Venn diagrams to identify potential lysosomal TMEM106B-binding proteins. Total 848 proteins were identified in immunoprecipitates from WT brains but 268 of them were also found in those from TMEM106B-deficient brains. Thus, 580 proteins were considered as potential TMEM106B-binding proteins. Among them, 22 proteins were lysosomal proteins based on The Mouse Lysosomal Gene Database (http://lysosome.unipg.it/mouse.php). **d)** Biosynthesis and degradation of GalCer and ST. SMase, sphingomyelinase; SMS, sphingomyelin synthase; CGT, cerebroside galactosyl transferase; CST, cerebroside sulfotransferase; ASA, arylsulfatase A. TMEM106B deficiency results in a decrease in levels of GalCer and ST (blue) while having no effects on levels of CER and SM (green). **e)** Schematic drawing of full-length (FL) TMEM106B-GFP and TMEM106B-GFP lacking amino acids (aa)123–275 (ΔC) and aa6–94 (ΔN). **f)** Representative blots of co-IP assays using HEK293T cells expressing GFP, FL TMEM106B-GFP, ΔC TMEM106B-GFP, or ΔN TMEM106B-GFP, together with myc-DDK-tagged GALC. **g)** Quantification of co-IP in (**f**). Mean ± SEM, n = 3-4 experiments, **p < 0.01, ****p < 0.0001; one-way ANOVA with Dunnett’s post hoc test comparing each sample with FL TMEM106B-GFP.

LC-MS/MS analysis of elution of anti-TMEM106B immunoprecipitates identified 22 potential lysosomal TMEM106B-binding proteins, some of which were previously reported to interact with TMEM106B, such as adaptor protein subunits, vacuolar-ATPase V0 domain subunits, and cathepsin D (Feng et al., 2020; Klein et al., 2017; Stagi et al., 2014), further validating our immunoprecipitation (**Figure 3C**). Interestingly, one of the TMEM106B-binding proteins identified was galactosylceramidase (GALC), an enzyme that hydrolyzes GalCer (**Figure 3C and D**). Currently, there are no commercially available anti-GALC antibodies that reliably detect endogenous GALC in mouse brains. To confirm the results of MS analysis, we thus performed co-IP assay using HEK293T cells transiently expressing GALC and green fluorescent protein (GFP), full-length (FL) TMEM106B-GFP, ΔC-TMEM106B-GFP, or ΔN-TMEM106B-GFP. We found that GALC was significantly co-immunoprecipitated with FL TMEM106B-GFP or ΔN-TMEM106B-GFP, but not GFP or ΔC-TMEM106B-GFP (**Figure 3E-G**). These results suggest that TMEM106B interacts with GALC in the lysosomal lumen. T185S and D252N mutations in TMEM106B are associated with human FTLD and HLD, respectively (Feng *et al*., 2021). These mutations however have no significant effects on co-IP of GALC (**Supplementary Figure 1**).

To further examine whether TMEM106B regulates GALC activity, we performed GALC activity assay using several brain regions of WT and TMEM106B-deficient mice. GALC activity can be measured using the fluorogenic substrate 4-methylumbelliferyl β-D-galactopyranoside (MUGAL) in the presence of AgNO_3_, which specifically inhibits β-galactosidase (β-Gal) (Martino et al., 2009). We found that GALC activity was significantly increased in the forebrain and brainstem but not cerebellum of TMEM106B-deficient mice (**Figure 4**). Together, these results suggest that TMEM106B binds to GALC and regulates GALC activity in most but not all brain regions.

**Figure 4:**
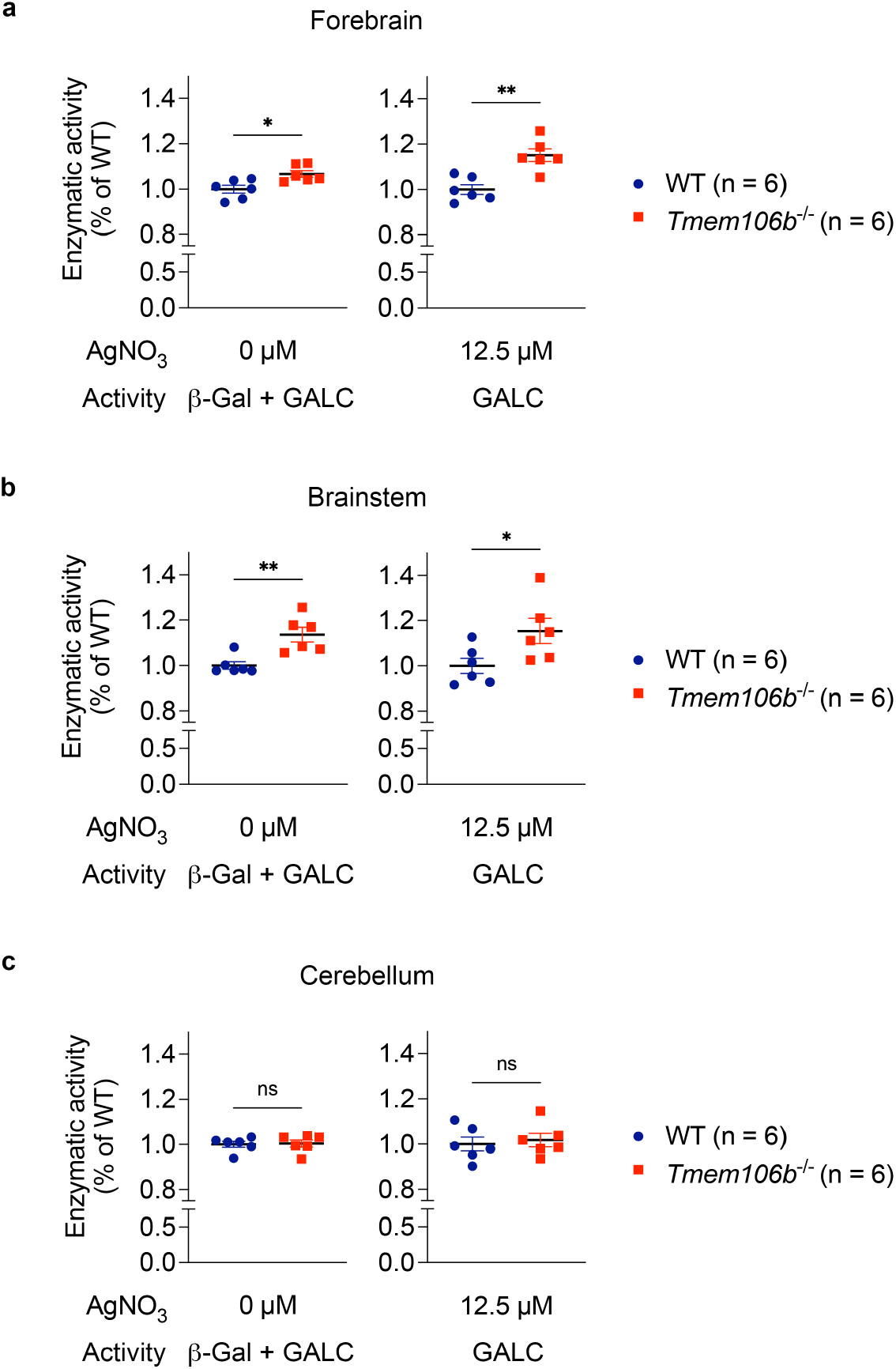
TMEM106B regulates GALC activity in the brain. **a)** Enzymatic activity of forebrain extracts of WT and *Tmem106b*^−/−^ mice at 13 months of age toward MUGAL substrate in the absent or presence of 12.5 μM AgNO_3_. n = 6 mice (2 males and 4 females) per genotype. *p < 0.05, **p < 0.01; Two-tailed unpaired t-test. **b)** Enzymatic activity of brainstem extracts of WT and *Tmem106b*^−/−^ mice at 13 months of age toward MUGAL substrate in the absence or presence of 12.5 μM AgNO_3_. n = 6 mice (2 males and 4 females) per genotype. *p < 0.05, **p < 0.01; Two-tailed unpaired t-test. **c)** Enzymatic activity of brainstem extracts of WT and *Tmem106b*^−/−^ mice at 13 months of age toward MUGAL substrate in the absence or presence of 12.5 μM AgNO_3_. n = 6 (2 males and d) 4 females) mice per genotype. Regardless of the absence or presence of AgNO_3_, two genotypes were not significantly different by two-tailed unpaired t-test.

## Discussion

The major finding of the present study is that endolysosomal protein TMEM106B regulates myelin lipid metabolism. Our shotgun lipidomics and subsequent SFC-MS/MS revealed that levels of two major classes of myelin lipids GalCer and ST, which together constitute ∼27% of the myelin lipid (Marcus and Popko, 2002), were significantly decreased in the brain of TMEM106B-deficient mice. Our results could potentially explain how a mutation in lysosomal TMEM106B leads to HLD. Importantly, dysfunction of myelin and/or myelin lipid metabolism has also been reported in other TMEM106B-associated brain disorders such as FTLD and AD. A recent study has shown disrupted myelin lipid metabolism in FTLD-*GRN* patients (Marian *et al*., 2023). White matter abnormality has long been known in AD (Bartzokis et al., 2003; Brun and Englund, 1986; Englund, 1998; Migliaccio et al., 2012; Nasrabady et al., 2018; Stout et al., 1996). In addition, several studies have reported a significant reduction in ST levels in preclinical AD cases (Cheng et al., 2013; Han et al., 2002; Han et al., 2003; Wallin et al., 1989). Interestingly, a previous rodent study has revealed that adult-onset ST deficiency is sufficient to cause AD-like neuroinflammation and cognitive impairment (Qiu et al., 2021). ST has also been shown to bind with high affinity to TREM2, a microglial AD risk factor (Wang et al., 2015). These studies raise an intriguing possibility that the TMEM106B variant may be involved in the brain disorders by altering levels of GalCer and ST. Outside the brain, TMEM106B has been identified as a driver of lung cancer metastasis (Grzeskowiak *et al*., 2018; Kundu *et al*., 2018) and as a proviral host factor for SARS-CoV-2, a coronavirus that caused the COVID19 pandemic (Baggen *et al*., 2021; Schneider *et al*., 2021; Wang *et al*., 2021). It will be interesting to examine whether a reduction in GalCer or ST is also observed in other TMEM106B-deficient tissues and protects against the lung cancer metastasis or SARS-CoV-2 infection.

In addition to GalCer and ST, TMEM106B deficiency has been shown to affect levels of other lipids such as PA and several species of PC and PE in our lipidomic analysis. How TMEM106B deficiency affects other lipids and whether those lipids are involved in TMEM106B-associated diseases remain unclear. Several previous studies have reported that PGRN deficiency results in alteration in the levels of BMP, glucosylshingosine (GlcSph), GlcCer, and/or gangliosides in human and/or mouse models of FTLD-*GRN* (Boland *et al*., 2022; Logan *et al*., 2021). However, these lipids were not significantly altered in TMEM106B-deficient brains, although TMEM106B is a risk modifier of FTLD-*GRN*. Further work will be required to better understand our lipidomic results and link them to previous findings.

Our unbiased IP and LC-MS/MS analysis identified GALC, an enzyme that hydrolyzes GalCer, as a novel interaction partner for TMEM106B. It appears that TMEM106B binds and regulates activity of GALC in the lysosomal lumen, which explains reduced levels of both GalCer and ST in TMEM106B-deficient brains as ST is produced by sulfation of GalCer through the action of cerebroside sulfotransferase (Blomqvist et al., 2021). Although the T185 variant and D252N mutation in TMEM106B have been associated with FTLD and HLD, respectively (Feng *et al*., 2021), our co-IP assay showed no significant effects of the variant and mutation on TMEM106B binding to GALC. Whether the negative result is due to our assay system overexpressing both TMEM106B and GALC or whether the genetic variant and mutation influence activity of GALC rather than TMEM106B binding to GALC is currently unclear and requires further investigation. Recent cryo-electron microscopy studies found amyloid fibrils composed of C-terminal TMEM106B in human brains (Chang *et al*., 2022; Fan *et al*., 2022; Jiang *et al*., 2022; Schweighauser *et al*., 2022). Our results suggest that the C-terminal fragment of TMEM106B could also interact with GALC. Effects of the C-terminal fragment on GALC activity in the brain will be an important avenue for future studies.

In conclusion, accumulating evidence clearly suggests that lysosomal TMEM106B plays a key role in a wide range of human conditions. Our study revealed a novel function of TMEM106B regulating myelin lipid metabolism by physically interacting with GALC. These findings provide mechanistic insights into TMEM106B-associated disorders and offer potential therapeutic strategies targeting TMEM106B-GALC interaction or modulating GalCer/ST levels.

## Author Contributions

Conceptualization, H.T. and S.M.S.; Methodology, H.T., X.H., and S.M.S.; Investigation, H.T., A.P.-C. H.Y. and X.H.; Writing – Original Draft, H.T. and S.M.S.; Writing – Review & Editing, all; Funding Acquisition, S.M.S.; Supervision, S.M.S.

## Acknowledgments

We thank Kristin DeLuca for assistance with mouse husbandry. This work was supported by National Institute on Aging of the National Institutes of Health under grant number R01AG034924 and R01AG066165 to S.M.S. We also thank the MS & Proteomics Resource at Yale University for providing the necessary mass spectrometers and the accompany biotechnology tools funded in part by the Yale School of Medicine and by the Office of The Director, National Institutes of Health (S10OD02365101A1, S10OD019967, and S10OD018034). This work was supported in part by the Lipidomics Shared Resource, Hollings Cancer Center, Medical University of South Carolina (P30 CA138313 and P30 GM103339). This work was also supported by a grant from National Institute on Aging (RF1 AG061729) to X.H. The content is solely the responsibility of the authors and does not necessarily represent the official views of the National Institutes of Health.

## Methods

### Lead contact

Further information and requests for resources and reagents should be directed to and will be fulfilled by the Lead Contact, Stephen M. Strittmatter.

## Data and code availability

LC-MS/MS raw data for Figure 3 have been deposited at PRIDE and are publicly available as of the date of publication. Accession numbers will be described. This paper does not report any codes. Any additional information required to reanalyze the data reported in this paper is available from the lead contact upon request.

### Mouse strains

TMEM106B-deficient mice on C57BL/6 background were described previously (Klein *et al*., 2017). This *Tmem106b*^−/−^ line was generated by LacZ gene trap strategy and expresses 5 to 10% residual full-length TMEM106B protein (Perez-Canamas et al., 2021; Zhou et al., 2020). All mice were housed in groups of 2 to 5 animals per cage with *ad libitum* access to standard mouse chow and water. Lighting was maintained at 12:12 light: dark cycle. All animal protocols were approved by the Yale Institutional Animal Care and Use Committee. Both male and female were used in this study and the numbers are described in Figure legends.

### Mouse brain tissue preparation

Mice were euthanized using CO_2_. After transcardiac perfusion using ice-cold DPBS, the brains were removed, and the hemispheres were divided and stored at -80°C until use. For GALC activity assay, forebrain and hindbrain were separated and the hindbrain were further dissected into cerebellum and brainstem. The dissected brain regions were snap-frozen in liquid nitrogen and stored at -80°C until use.

### Lipidomic analysis

Lipid species were analyzed using multidimensional mass spectrometry-based shotgun lipidomic analysis as previously described (Han, 2016). In brief, the brain hemispheres from 12-month-old TMEM106B-deficient mice and WT littermates were homogenized in 0.1X PBS and the brain homogenate containing 1.0 mg of protein, which was determined with a Pierce™ BCA protein assay kit (ThermoFisher SCIENTIFIC #23225), was accurately transferred to a disposable glass culture test tube. A premixture of lipid internal standards was added prior to conducting lipid extraction for quantification of the targeted lipid species. Lipid extraction was performed using a modified Bligh and Dyer procedure (Wang and Han, 2014), and each lipid extract was reconstituted in chloroform:methanol (1:1, *v:v*) at a volume of 400 μL/mg protein. Phosphatidylethanolamine (PE) and cholesterol (CHL) were derivatized as described previously (Cheng et al., 2007; Han et al., 2005) before lipidomic analysis.

For shotgun lipidomics, lipid extract was further diluted to a final concentration of ∼500 fmol total lipids per μL. Mass spectrometric analysis was performed on a triple quadrupole mass spectrometer (TSQ Altis, Thermo Fisher Scientific, San Jose, CA) and a Q Exactive mass spectrometer (Thermo Scientific, San Jose, CA), both of which were equipped with an automated nanospray device (TriVersa NanoMate, Advion Bioscience Ltd., Ithaca, NY) as described (Han et al., 2008). Identification and quantification of lipid species were performed using an automated software program (Wang et al., 2016; Yang et al., 2009). Data processing (e.g., ion peak selection, baseline correction, data transfer, peak intensity comparison and quantitation) was performed as described (Yang *et al*., 2009). The result was normalized to the protein content (nmol lipid/mg protein).

### GluCer and GalCer species analysis separated by SFC-MS/MS

Separation of GlcCer and GalCer species was performed at the Lipidomics Shared Resource at Medical University of South Carolina. Lipids were extracted from the 0.1X PBS homogenate containing 1 mg of protein, and levels of GlcCer and GalCer species were measured with supercritical fluid chromatography-tandem mass spectrometry (SFC-MS/MS) analysis.

### Immunoprecipitation

Immunoprecipitation using mouse brain lysates was performed as previously reported (Perez-Canamas *et al*., 2021). WT and *Tmem106b*^-/-^ brains at 6-7-months of age were homogenized in ice-cold TBST (50 mM Tris-HCl, pH 7.4, 150 mM NaCl, 1% Triton X-100) supplemented with cOmplete-mini (Roche). After ultracentrifugation at 100,000 x g, the supernatant was pre-cleared using Protein A-Sepharose CL-4B (SIGMA #17-0780-01) for 3 h at 4°C. Then, anti-TMEM106B antibody (Abcam #ab140185) covalently linked to Protein A-Sepharose CL-4B with BS^3^ (ThermoScientific #21580) was added to the pre-cleared lysates. After overnight incubation at 4°C, the immunoprecipitates were washed six times with ice-cold TBST and proteins were eluted with 2 x Laemmli buffer (Bio-Rad).

### In Solution Protein Digestion

A methanol-chloroform precipitation was performed according to standard protocols. The protein pellet was resuspended in 20 μL 8M urea, 0.4 M ammonium bicarbonate with vigorous vortexing. Proteins were reduced by the addition of 2.0 μL 45 mM dithiothreitol (ThermoFisher SCIENTIFIC #20290) and incubation at 37ºC for 30 min, and then alkylated with the addition of 2.0 μL 100 mM iodoacetamide (SIGMA #I1149) and incubation in the dark at room temperature for 30 min. The urea was diluted to 2 M by adding 55 μL of water. Samples were then enzymatically digested using 1.0 μL 0.5 mg/ml trypsin (Promega Seq. Grade Mod. Trypsin, # V5113) and incubation at 37ºC for 16 h. Digestion was halted by acidification with trifluoroacetic acid (TFA) to 0.1%. The mixtures were desalted using C18 Ultra microspin columns (The Nest Group, #SUM SS18V) following the manufacturer’s directions. Peptides were eluted with 2 x 160 μL 0.1% TFA, 80% acetonitrile, then speedvaced dry. Peptides were dissolved in 32 μL MS loading buffer (2% aceotonitrile, 0.2% trifluoroacetic acid). A nanodrop measurement (Thermo Scientific Nanodrop 2000 UV-Vis Spectrophotometer) determined protein concentrations (A260/A280). Each sample was then further diluted with MS loading buffer to 0.04 μg/μL, with 200 ng (5 μL) injected for LC-MS/MS analysis.

### LC-MS/MS on the Thermo Scientific Q Exactive Plus

LC-MS/MS analysis was performed on a Thermo Scientific Q Exactive Plus equipped with a Waters nanoAcquity UPLC system utilizing a binary solvent system (A: 100% water, 0.1% formic acid; B: 100% acetonitrile, 0.1% formic acid). Trapping was performed at 5 μL/min, 97% Buffer A for 3 min using a Waters Symmetry® C18 180 μm x 20 mm trap column. Peptides were separated using an ACQUITY UPLC PST (BEH) C18 nanoACQUITY Column 1.7 μm, 75 μm x 250 mm (37ºC) and eluted at 300 nL/min with the following gradient: 3% buffer B at initial conditions; 5% B at 1 min; 35% B at 90 min; 50% B at 105 min; 90% B at 110 min, 90% B at 115 min; return to initial conditions at 116 min. MS was acquired in profile mode over the 300-1,700 m/z range using 1 microscan, 70,000 resolution, AGC target of 3E6, and a maximum injection time of 45 ms. Data dependent MS/MS were acquired in centroid mode on the top 20 precursors per MS scan using 1 microscan, 17,500 resolution, AGC target of 1E5, maximum injection time of 100 ms, and an isolation window of 1.7 m/z. Precursors were fragmented by HCD activation with a collision energy of 28%. MS/MS were collected on species with an intensity threshold of 2E4, charge states 2-6, and peptide match preferred. Dynamic exclusion was set to 20 s.

### Peptide Identification

Data was analyzed using Proteome Discoverer (version 1.3) software and searched in-house using the Mascot algorithm (version 2.6.0) (Matrix Science). The data was searched against a SwissProtein database with taxonomy restricted to Mus musculus. Search parameters used were trypsin digestion with up to 2 missed cleavages; peptide mass tolerance of 10 ppm; MS/MS fragment tolerance of 0.02 Da; and variable modifications of methionine oxidation and carbamidomethyl cysteine. Normal and decoy database searches were run, with the confidence level was set to 95% (p<0.05).

### Co-immunoprecipitation (co-IP) using HEK293T cells

Co-IP was performed as previously reported (Klein *et al*., 2017) with slight modifications. Plasmids encoding myc-DDK-tagged GALC (#MR225749) were purchased from OriGene Technologies, Inc. Plasmids encoding human TMEM106B-mCherry was generated using pmCherry-N1 vector (Clontech #632523). T185S and D252N mutations were introduced using Quik change II XL site-directed mutagenesis kit (Agilent #200521). Two days after transfection using Lipofectamine™ 2000 (ThermoFisher SCIENTIFIC #11668019), HEK293T cells were harvested and lysed with ice-cold 50 mM Tris-HCl pH 7.5, 150 mM NaCl, 1% Triton X-100 supplemented with cOmplete-mini and PhosSTOP (Roche). After ultracentrifugation at 100,000 x g for 30 min, the supernatant was incubated with ChromoTek GFP-or RFP-trap® Agarose (proteintech #gta or #rta) for 3 h at 4°C. The immunoprecipitates were washed five times with ice-cold lysis buffer and boiled with 2 x Laemmli Buffer (Bio-Rad) with βME. Immunoblot was performed as previously reported (Klein *et al*., 2017).

### GALC activity assay

GALC activity assay was performed as previously reported with modifications (Martino *et al*., 2009). Briefly, frozen dissected brain tissues (forebrain, brainstem, and cerebellum) were homogenized using a dounce homogenizer in citrate/phosphate (CP) buffer pH 5.2, 1% Triton X-100, supplemented with cOmplete-mini and phosSTOP (Roche) on ice and incubated for 30 min at 4°C. The supernatants after ultracentrifugation at 100,000 x g were used for GALC activity assay. Protein concentrations were determined by Pierce™ BCA protein assay kit (ThermoFisher SCIENTIFIC #23225) and used for normalization. Five micro litters of the supernatants were mixed and preincubated with 10 μL of H_2_O (vehicle) or AgNO_3_ (SIGMA #204390) (final concentration 12.5 μM) in CP buffer pH 4 in 96-well black plate (Costar #3916) for 10 min at room temperature (RT) and then added 30 μL of 4-methylumbelliferyl β-D-galactopyranoside (MUGAL) (final concentration 0.8 mM) (SIGMA #M1633) and incubated for 1 h at 37°C. Reaction was stopped by adding 200 μL of 0.5M glycine-NaOH pH 10.5. Measurements were taken with VICTOR Nivo® multimode plate reader (PerkinElmer) at excitation of 355 nm and emission of 460 nm.

## Statistical analysis

Two-tailed unpaired t test (for 2 groups) and one-way ANOVA (for > 3 groups) were performed using GraphPad Prism (version 9.2.0). All data are shown as mean ± SEM and specific n values are reported in each Figure legend. Data are considered to be statistically significant if p < 0.05.

## Figures

**Supplementary Figure 1:**
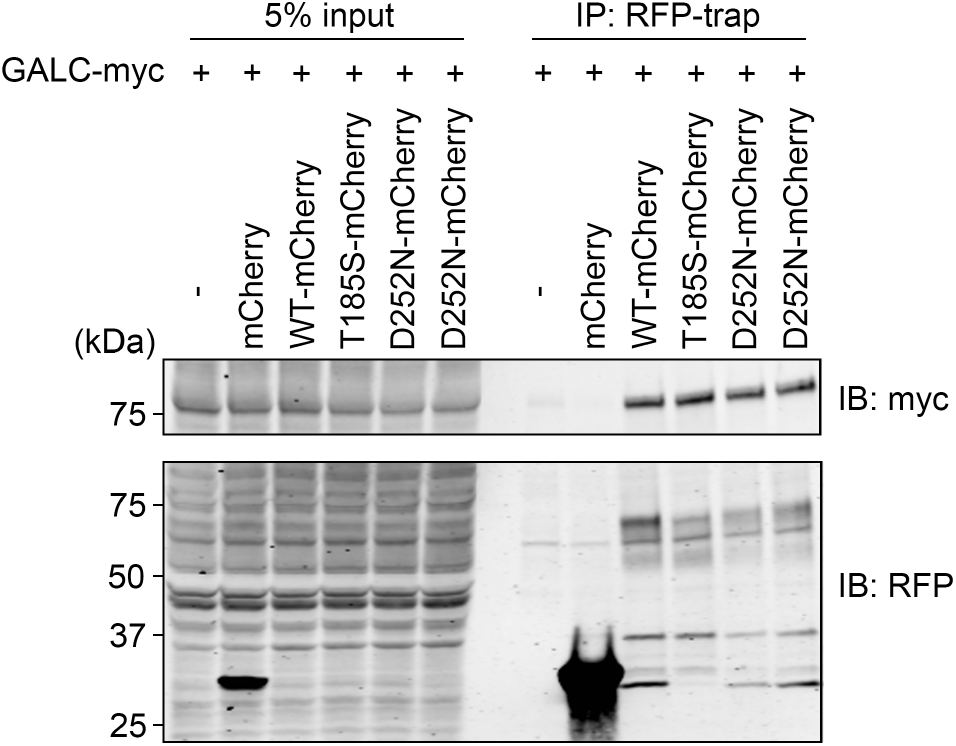
T185S and D252N mutations have no significant effects on TMEM106B binding to GALC. Representative blots of co-IP assays using HEK293T cells expressing mCherry, FL TMEM106B-mCherry, T185S TMEM106B-mCherry, or D252N TMEM106B-mCherry, together with myc-tagged GALC.

